# Genome-enabled insights into the ecophysiology of the comammox bacterium *Candidatus* Nitrospira nitrosa

**DOI:** 10.1101/144600

**Authors:** Pamela Y. Camejo, Jorge Santo Domingo, Katherine D. McMahon, Daniel R. Noguera

**Affiliations:** Department of Civil and Environmental Engineering, University of Wisconsin - Madison, Madison, WI, USA; Environmental Protection Agency, Cincinnati, OH, USA; Department of Bacteriology, University of Wisconsin - Madison, Madison, WI, USA

**Author notes:** Email addresses. Corresponding author: Daniel R. Noguera, 1415 Engineering Drive, Madison, WI 53706.

## Abstract

The recently discovered comammox bacteria have the potential to completely oxidize ammonia to nitrate. These microorganisms are part of the *Nitrospira* genus and are present in a variety of environments, including Biological Nutrient Removal (BNR) systems. However, the physiological traits within and between comammox- and nitrite oxidizing bacteria (NOB)-like *Nitrospira* species have not been analyzed in these ecosystems. In this study, we identified *Nitrospira* strains dominating the nitrifying community of a sequencing batch reactor (SBR) performing BNR under micro-aerobic conditions. We recovered metagenomes-derived draft genomes from two *Nitrospira* strains: (1) *Nitrospira* sp. UW-LDO-01, a comammox-like organism classified as *Candidatus* Nitrospira nitrosa, and (2) *Nitrospira* sp. UW-LDO-02, a nitrite oxidizing strain belonging to the *Nitrospira defluvii* species. A comparative genomic analysis of these strains with other *Nitrospira-like* genomes identified genomic differences in *Ca.* Nitrospira nitrosa mainly attributed to each strains’ niche adaptation. Traits associated with energy metabolism also differentiate comammox from NOB-like genomes. We also identified several transcriptionally regulated adaptive traits, including stress tolerance, biofilm formation and micro-aerobic metabolism, which might explain survival of *Nitrospira* under multiple environmental conditions. Overall, our analysis expanded our understanding of the genetic functional features of *Ca.* Nitrospira nitrosa, and identified genomic traits that further illuminate the phylogenetic diversity and metabolic plasticity of the *Nitrospira* genus.

## INTRODUCTION

Nitrification is a microbiological process that plays an important role in the nitrogen (N) cycle. This process has been conventionally known as a two-step reaction. The first step, oxidation of ammonia to nitrite, is performed by ammonia-oxidizing bacteria (AOB) or archaea (AOA) and the second step, oxidation of nitrite to nitrate, is carried out by nitrite-oxidizing bacteria (NOB). Recently, the discovery of a new player with the potential to completely oxidize ammonia to nitrate, as in the case of complete ammonia oxidizing (comammox) organisms,^1, 2^ has dramatically changed our understanding of microbial mediated N transformations in engineered and natural systems.

Comammox bacteria have been classified within the genus *Nitrospira.* Members of this genus were conventionally regarded as NOB and were thought to rely only on nitrite for growth. However, the genomes of the four comammox-like *Nitrospira* identified to date (*Ca.* Nitrospira nitrosa, *Ca.* Nitrospira nitrificans, *Ca.* Nitrospira inopinata and *Nitrospira* sp. Ga0074138^1-3^), encode the genes necessary for ammonia and nitrite oxidation, suggesting that *Nitrospira* are much more metabolically versatile organisms. Furthermore, comammox-like *Nitrospira* have been identified in a variety of habitats, including groundwater wells, drinking water biofilters, wastewater treatment plants (WWTPs) and other soil and aquatic environments.^4^ These findings have prompted questions regarding the ecological significance and lifestyle of these organisms in each of these ecosystems.

Nutrient removal in WWTPs relies on nitrifying organisms to remove N from the wastewater. *Nitrospira-like* bacteria appear to be the dominant nitrite-oxidizers^5-7^ in most WWTPs and laboratory scale reactors. The abundance of comammox in WWTPs has been briefly surveyed, and preliminary results show this functional group is present in these systems.^4^ However, genetic and functional adaptations of comammox to this environment have not been addressed.

In this study, the community performing N removal in a Biological Nutrient Removal (BNR) lab-scale reactor was analyzed to explore the genomic basis for comammox ecophysiology. A sequencing batch reactor (SBR) was operated under cyclic anaerobic and micro-aerobic conditions using two different operational conditions. During the first stage (nitrite addition during a micro-aerobic phase), two *Nitrospira-like* strains were enriched in the reactor. Draft genome sequences of these two strains were assembled from metagenomic data; one of them was identified as a commamox organism, the other as an NOB. Here, we used the draft genomes of these strains, as well as genomes from both NOB- and comammox-related bacteria, to perform a comparative genome analysis of the genus *Nitrospira*.

## MATERIAL AND METHODS

### Operation of Lab-Scale Sequencing Batch Reactor

A laboratory-scale SBR was originally inoculated with activated sludge obtained from the Nine Springs WWTP in Madison, WI, which uses a modified University of Cape Town (UCT) process designed to achieve biological P removal^8^ and operates with high aeration rates.^9^ Synthetic wastewater containing acetate as the sole carbon source was used for the feed, as described elsewhere.^10^ The hydraulic retention time (HRT) and solids retention time (SRT) were 24 h and 80 days, respectively. The pH in the system was controlled to be between 7.0 and 7.5.

The 2-liter reactor was operated under alternating anaerobic and low oxygen cycles. During Stage 1 of operation the cycles consisted of 2 h anaerobic, 5 h micro-aerobic, 50 min settling and 10 min decanting. At the beginning of the micro-aerobic phase, sodium nitrite was added to reach an in-reactor concentration of 10 mg N-N0_2_^-^/L and stimulate the use of nitrite as an electron donor. In addition, an on/off control system was used to limit the amount of oxygen pumped to the reactor (0.02 L/min) and maintain low dissolved oxygen (DO) concentrations in the mixed liquor, as described elsewhere.^10^ After 100 days of operation, the nitrite supplement was eliminated and the reactor cycle was changed to: 1.5 h anaerobic, 5.5 h micro-aerobic, 50 min settling and 10 min decanting (Stage 2).

### Sample Collection and Analytical Tests

To monitor reactor performance, mixed liquor and effluent samples were collected, filtered through a membrane filter (0.45 μm; Whatman, Maidstone, UK) and analyzed for acetate, PO_4_^3-^-P, NH_4_+-N, NO_3_^-^-N, and NO_2_^-^-N. The concentrations of PO_4_^3-^−P were determined according to Standard Methods.^11^ Total ammonia (NH_3_ + NH_4_+) concentrations were analyzed using the salicylate method (Method 10031, Hach Company, Loveland, CO). Acetate, nitrite and nitrate were measured using high-pressure liquid chromatography as previously described.^10^

Biomass samples from the reactors were collected weekly and stored at -80°C until DNA extraction was performed. DNA was extracted using UltraClean^®^ Soil DNA Isolation Kit (MoBIO Laboratories, Carlsbad, CA). Extracted DNA was quantified using a NanoDrop spectrophotometer (Thermo Fisher Scientific, Waltham, MA) and stored at -80°C.

### Metagenome Sequencing, Assembly and Binning

Samples from day 100 (Stage 1) and days 317, 522 and 674 (Stage 2) were selected for metagenomic analysis. Illumina TruSeq DNA PCR free libraries were prepared for DNA extracts according to the manufacturer’s protocol and paired-end sequenced on either the Illumina HiSeq 2000 platform (v4 chemistry, 2 × 150 bp; 522-day sample), or the Illumina MiSeq platform (v3 chemistry, 2 × 250 bp; other samples). Unmerged reads were quality-trimmed and filtered with Sickle (https://github.com/ucdavis-bioinformatics/sickle.git), using a minimum phred score of 20 and a minimum length of 50 bp. The metagenomic reads from the 100-day sample (Stage 1) were assembled using IDBA-UD.^12^ Individual genome bins were extracted from the metagenome assembly with the R package ‘mmgenome’^13^ using the differential coverage principle.^14^ The bins were initially extracted by plotting the genome coverage of contigs in metagenomes from day 100 and 317. During the bin extraction, GC content and taxonomy of contigs were also taken into consideration.

After binning, SSPACE was used to filter small scaffolds (length < 1,000 bp), extend scaffolds, and to fill gaps.^15^ Genome completeness and contamination was estimated using CHECKM 0.7.1.^16^ Table S1 displays quality metrics of the draft genomes after each of the steps previously described. Two putative *Nitrospira-like* bins were identified and annotated using MetaPathways v 2.0.^17^ To further reduce contamination in these assembled bins, scaffolds with Open Reading Frames (ORFs) having less than 85% nucleotide identity and 0% protein identity to other *Nitrospira* genomes were removed from the bins.

### Average Nucleotide Identity (ANI)

Pair-wise ANI values of *Nitrospira-like* genomes were obtained using the ANIm method^18^ and implemented in the Python script ‘calculate_ani.py’ available at https://github.com/ctSkennerton/scriptShed/blob/master/calculate_ani.py.

### Phylogenetic Analyses

The phylogeny of the draft genomes was assessed by constructing a phylogenetic tree using a concatenated alignment of marker genes. First, PhyloSift v 1.0.1^19^ was used to extract a set of 38 marker genes from each genome. Then, the extracted marker protein sequences were concatenated into a continuous alignment to construct a maximum-likelihood (ML) tree, using RAxML v 7.2.8.^20^ RAxML generated 100 rapid bootstrap replicates followed by a search for the best-scoring ML tree.

For phylogenetic analyses of ammonia monooxygenase subunit A *(amoA),* hydroxylamine reductase *(hao)* and nitrite oxido-reductase subunit A (nxrA) genes, full nucleotide datasets were downloaded from the NCBI GenBank database.^21^ Alignment was performed on sequences retrieved from the NCBI and the draft genomes using the ‘AlignSeqs’ command in the DECIPHER “R” package.^22^ Phylogenetic trees were calculated using neighbor-joining criterion with 1,000 bootstrap tests for every node, using the MEGA6 software package.^23^ Trees were visualized with the assistance of TreeGraph.^24^

### Population Structure by Metagenomic Analysis

To estimate the abundance of currently known ammonia oxidizers, comammox, and nitrite oxidizers in the reactor over time, paired-end DNA reads from the metagenomic datasets (days 100, 317, 522 and 674) were competitively mapped to the published genome sequence of 14 AOB *(Nitrosomonas, Nitrosospira* and *Nitrosococcus* genera), 6 AOA *(Nitrososphaera, Nitrosoarchaeum* and *Nitrosopumilus),* 5 NOB *(Nitrospira* and *Nitrobacter* lineage), 5 anaerobic ammonia-oxidizing (anammox) bacteria *(Ca.* Brocadia fulgida, *Ca.* Brocadia caroliniensis, *Ca.* Kuenenia stuttgartiensis, *Ca.* Brocadia sinica and *Ca.* Jettenia caeni), 4 comammox bacteria (Ca. Nitrospira nitrosa, *Ca.* Nitrospira nitrificans, *Ca.* Nitrospira inopinata and *Nitrospira* sp. Ga0074138) and the two *Nitrospira-like* draft genomes retrieved from the reactor, using the software package BBMap version 35.85 (https://sourceforge.net/projects/bbmap). A list of the genomes included in this analysis and the number of reads mapping to each sequence is found in Table S2. For each organism, the number of unambiguous reads (best hit) mapping to the genomic sequence with a minimum alignment identity of 90% was quantified and normalized by the number of reads in each metagenome, paired-end reads average length and genome size (Table S2).

### Orthologous Genes Clusters

To assess the degree of homology in the proteomes of the two *Nitrospira-like* genomes, orthologous genes clusters (OCs) were determined using OrthoMCL.^25^ OrthoMCL was run with a BLAST E-value cut-off of 1e-5, and an inflation parameter of 1.5.

Protein products of each ortholog set were compared against the Eggnog database^26^ using the eggNOG-mapper tool (https://github.com/jhcepas/eggnog-mapper) (*e*-value <10e^−2^) to determine the COG functional and super-functional category to which they belong.

### Accession Numbers

Raw reads and draft genome sequences have been submitted to NCBI and are accessible under the BioProject identifier PRJNA322674.

## RESULTS AND DISCUSSION

### Nutrient Removal in Lab-Scale Reactor

Results from a typical cycle of the lab-scale SBR at steady-state operation during Stage 1 are shown in Fig. 1. During the anaerobic phase, acetate added at the beginning of the anaerobic phase was completely consumed within an hour (Fig. 1B). P release to the mixed liquor during this condition was not observed (Fig. 1B), indicating the absence of polyphosphate accumulating organisms (PAO) in the reactor. Denitrification was incomplete, with only ~ 60% of the nitrate removed in the anaerobic phase (Fig. 1A), even though the reactor received acetate in this phase. This suggests that efficient acetate uptake was likely performed by glycogen accumulating organisms (GAO), without affecting P concentrations.^27^ In addition, nitrite production during this phase (~ 10%) is an indicator of partial denitrification (Fig. 1A).

**Figure 1.**
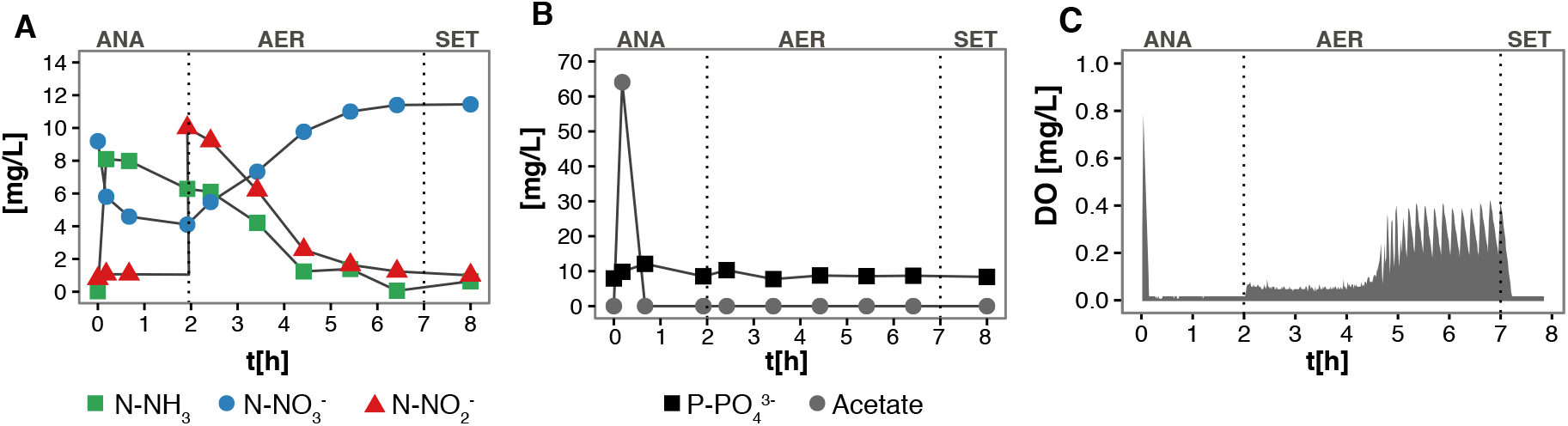
Nutrient profiles of (A) nitrogenous compounds, (B) phosphorous and acetate, and (C) oxygen concentration in a regular cycle of the lab-scale SBR during Stage 1. ANA: Anaerobic, AER: Micro-aerobic, SET: Settling.

Complete nitrification occurred in the subsequent micro-aerobic stage, where 91% ± 4% of ammonia and 91% ± 8% of the added nitrite were removed (Fig. 1A), with nitrate accumulation accounting for 50% ± 2% of the oxidized nitrogen. During the period of active nitrification, the DO remained below 0.05 mg/L (Fig. 1C), as the oxygen supplied balanced the oxygen uptake rate. The oxygen uptake rate decreased after nitrification ceased, and correspondingly, DO increased. To maintain a low-DO environment, aeration was stopped when DO exceeded the set point (0.2 mg O_2_/L) and resumed when DO decreased below the set point. This operation effectively maintained DO below 0.4 mg/L (Fig. 1C).

After 100 days of reactor operation under these conditions, the operational parameters were changed to promote simultaneous nitrification, denitrification and P removal (Stage 2). Unlike Stage 1, no nitrite was added to the reactor during micro-aerobic conditions. During this second stage, acetate added at the beginning of the anaerobic phase was used by PAOs for P-cycling, and nitrite and nitrate produced by ammonia oxidization were used as electron acceptors by PAOs during micro-aerobiosis, achieving simultaneous removal of N and P. Results of this stage were described elsewhere.^10^

### *Nitrospira-like* Genome Binning

Using a combination of bidimensional coverage and tetranucleotide frequency, two *Nitrospira-like* draft genomes were assembled from a sample collected at the end of Stage 1. The two draft genomes *(Nitrospira* sp. UW-LDO-1 and UW-LDO-2) had 3.9 and 3.5 Mbp in total with average GC content of 54.9% and 59.2%, respectively (Table S1). The reconstructed genomes were assessed to be nearly complete (completeness ≧90%) with low contamination (≦5%), according to the presence of 43 single-copy reference genes (Table S1).

Since the composite genomes did not encode complete 16S rRNA genes, the average nucleotide sequence identity (ANI) between the draft genomes assembled herein and formerly published *Nitrospira*-like genomes, was used to determine whether UW-LDO-1 and UW-LDO-2 represented distinct species, as this method has been shown to correlate well with previously defined 16S rRNA gene species boundaries.^28^ The calculated ANI and fraction of alignment for the *Nitrospira* genomes (Fig. 2) showed that UW-LDO-01 is a representative of the *Ca. Nitrospira nitrosa* species (ANI >94%, Fraction Aligned = 74.9%), while UW-LDO-02 had the closest nucleotide identity to *Nitrospira defluvii* (ANI = 92.4%, Fraction Aligned = 72.4%). None of the other ANI values were greater than 88%, indicating that the two genomes were different from each other, and supports their classification as *Nitrospira nitrosa* UW-LDO-01 and *Nitrospira defluvii* UW-LDO-02, respectively.

**Figure 2.**
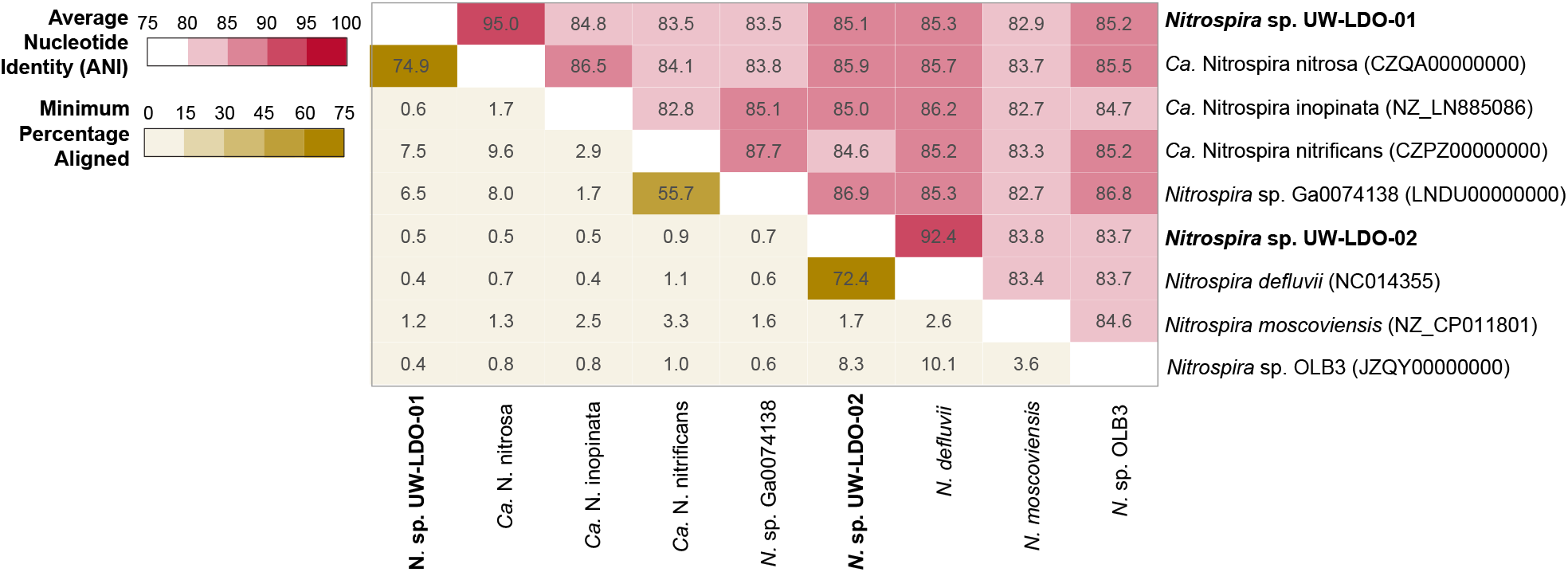
Comparison of the genome-wide average nucleotide identity and alignment percentage, of *Nitrospira-like* genomes. The heatmap shows the average nucleotide identity (red upper section of matrix) and the percentage of the two genomes that aligned (yellow lower section).

### Phylogenetic Analysis

A genome tree constructed from a concatenated protein alignment of 38 universally distributed single-copy marker genes^29^ confirms the affiliation of *Nitrospira* sp. UW-LDO-01 and UW-LDO-02 with *Ca.* Nitrospira nitrosa and *Nitrospira defluvii,* respectively (Fig. 3). Consistent with this phylogeny, UW-LDO-01 encoded the *amoCAB* operon, responsible for ammonia oxidation, while UW-LDO-02 did not.

**Figure 3.**
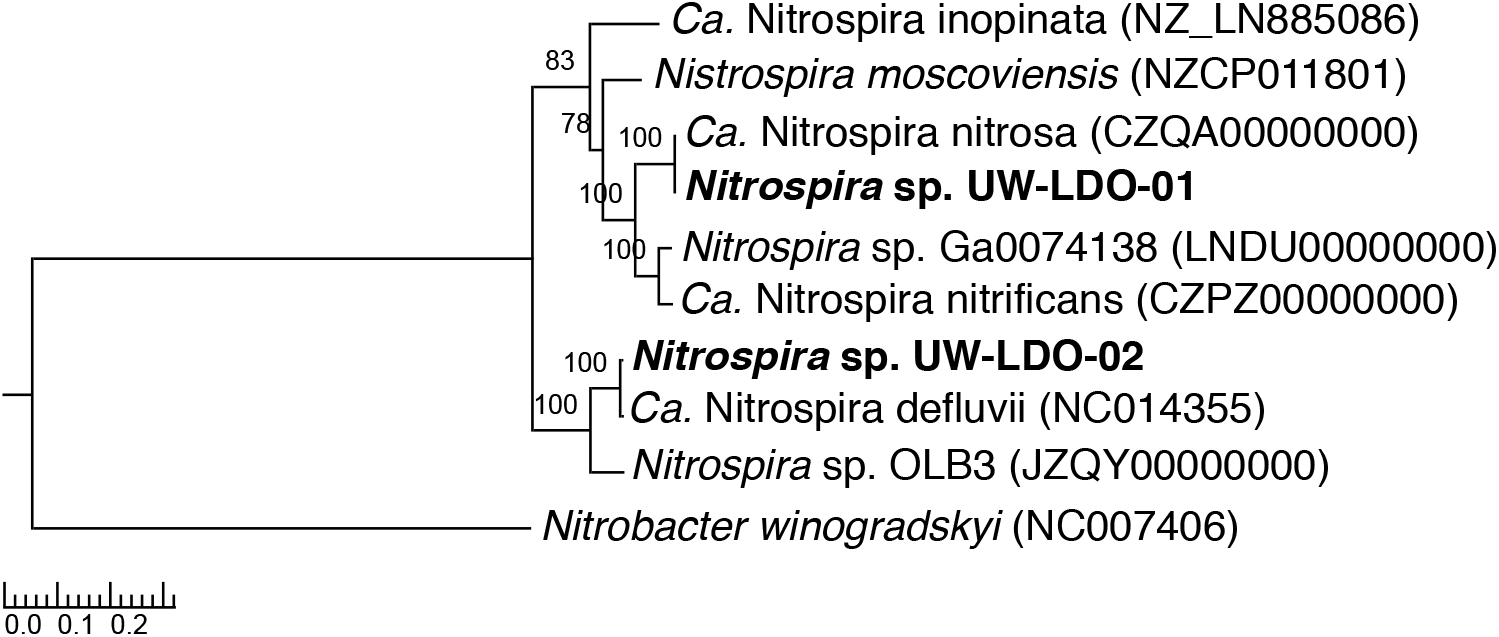
RAxML phylogenetic tree of a concatenated alignment of 37 marker genes (nucleotide sequence) data set with the root placed on the branch leading to *Nitrobacter winogradskyi.* The numbers at the nodes of both trees show support values derived from 100 RAxML bootstrap.

The *amoA* and *hao* genes are functional genes involved in redox nitrogen transformations and are also considered phylogenetic markers to study the diversity of ammonia oxidizing microorganisms (AOM).^30-33^ The phylogenetic tree topologies based on these genes (Fig. S1) further confirm the classification of UW-LDO-01 as related to *Ca.* Nitrospira nitrosa, although two paralogs of the *amoA* gene are present in the genome of *Ca. Nitrospira nitrosa,* while only one *amoA* gene was found in UW-LDO-1.

In addition, the *Nitrospira* sp. UW-LDO-1 and UW-LDO-2 genomes encoded the key enzyme for nitrite oxidation, *nxr,* which can also be used as a phylogenetic biomarker. UW-LDO-1 encoded two paralogs of the periplasmic NXR enzyme while *Nitrospira* UW-LDO-2 only encoded one copy. The affiliation of UW-LDO-1 with *Ca.* Nitrospira nitrosa and UW-LDO-2 with *N. defluvii* was supported by phylogeny based on the *nxrA* gene sequence (Fig. S2), consistent with the phylogenetic analysis of *amoA* and *hao.*

### Nitrifying Prokaryotes in Lab-Scale Reactor

The metagenomic analysis of the Stage 1 sample, which corresponds to the operational conditions when nitrite and ammonia were both present under micro-aerobic conditions, did not result in the assembly of any other genome of nitrifying microorganisms. Thus, in order to assess the relative abundance of other known nitrifying prokaryotes present in the reactor, we mapped metagenomic reads to published genomes of comammox, anammox, AOB, AOA and NOB, including *Nitrospira* sp. UW-LDO-01 and UW-LDO-02 (Fig. 3). After a competitive mapping of short-reads from metagenomic samples to each genome (>90% identity), the number of mapping reads was normalized to both metagenome size and reference genome size, and used as a proxy of genome abundance.

The metagenomic data show little evidence of AOA and anammox bacteria during the two stages (0.06% and 0.25% of mapping reads, respectively). AOB were detected in the system, albeit representing a small fraction of the community (0.17% and 0.05% of total number of reads during Stage 1 and 2, respectively). Noticeably, *Nitrospira-like* sequences (including comammox- and NOB-like genomes) recruited the greatest number of metagenomic reads (14.0% of total number of reads) in the Stage 1 sample (Fig. 4A). Within this genus, *Nitrospira sp.* UW-LDO-01 retrieved 32.3% of the reads competitively mapping to the *Nitrospira-like* genomes (Fig. 4B). The published genome of *Ca.* Nitrospira nitrosa retrieved 2.3% of the reads, while less than 4% mapped to other comammox genomes. Therefore, with only a small fraction of reads mapping to other ammonia oxidizers, we propose that *Nitrospira* sp. UW-LDO-01 was the main comammox in the reactor, and the main contributor to ammonia oxidation during Stage 1.

**Figure 4.**
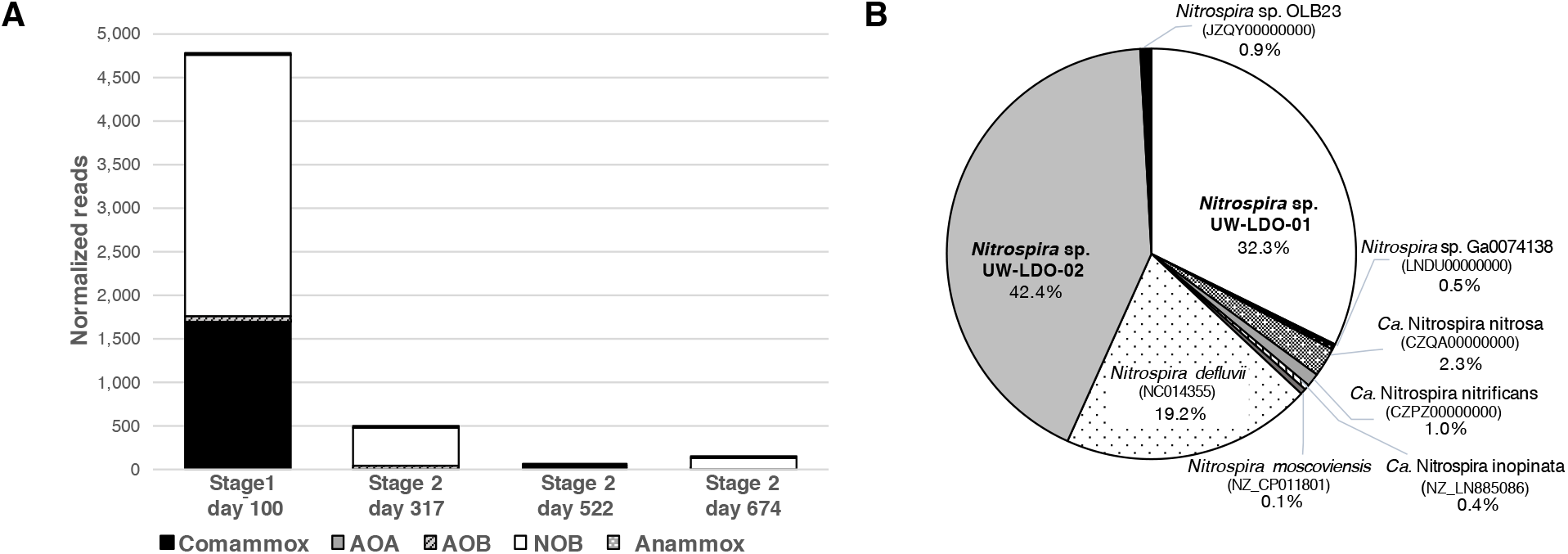
(A) Normalized frequency of metagenomic reads mapping to the genome of Comammox, AOB, AOA, NOB and Anammox-related organisms in samples from Stage 1 and 2 of the lab-scale SBR. (B) Relative abundance of reads mapping to genomes of *Nitrospira-related* bacteria in Stage 1 sample, including the draft genomes retrieved in this study.

*Nitrospira* sp. UW-LDO-02 appeared to be the most abundant NOB in the reactor during Stage 1, retrieving 42.4% of the *Nitrospira-like* reads (Fig. 4B), although a large fraction of reads competitively mapping to *N. defluvii* may indicate the presence of other nitrite oxidizing strains in the reactor. Therefore, the nitrite oxidation activity in the reactor was carried out by comammox and NOB.

The metagenomic analysis of Stage 2 samples reveals an overall decrease in the relative abundance of nitrifying organisms in the reactor after transitioning to this operational configuration (Fig. 4A). During this stage, metagenomic reads mapping to NOB and comammox genomes (including *Nitrospira* sp. UW-LDO-01 and UW-LDO-02) decreased to less than 1% of the total number of reads. This was in part due to the removal of nitrite addition during Stage 2. However, the decrease in comammox did not correspond to an increase in the number of reads mapping to other known AOM (Fig. 4A), suggesting the presence of still unrecognized AOM in reactors operated with low DO conditions, as previously reported.^34^

### Differential Gene Content Among *Ca.* Nitrospira nitrosa Genomes

Since *Nitrospira* sp. UW-LDO-01 is the second comammox genome representative of *Ca.* Nitrospira nitrosa, and the first comammox genome recovered from a nutrient removal bioreactor, a comparative analysis of its genetic content was carried out. First, a comparison of gene content among *Ca.* Nitrospira nitrosa (CZQA00000000) and *Nitrospira* sp. UW-LDO-01 was conducted by BlastP comparison of the translated coding DNA sequence (CDS) set, clustering of ortholog proteins, and annotation of representatives of each ortholog cluster (OC) and genome-unique CDSs.

Overall, sequencing and annotation of the UW-LDO-01 genome revealed a genomic inventory highly similar to the genome of *Ca.* Nitrospira nitrosa.^2^ The two genomes shared 81.2% of the OCs (3,202 OCs), with UW-LDO-01 and *Ca.* Nitrospira nitrosa having 740 and 857 unique OCs, respectively (Fig. 5A).

**Figure 5.**
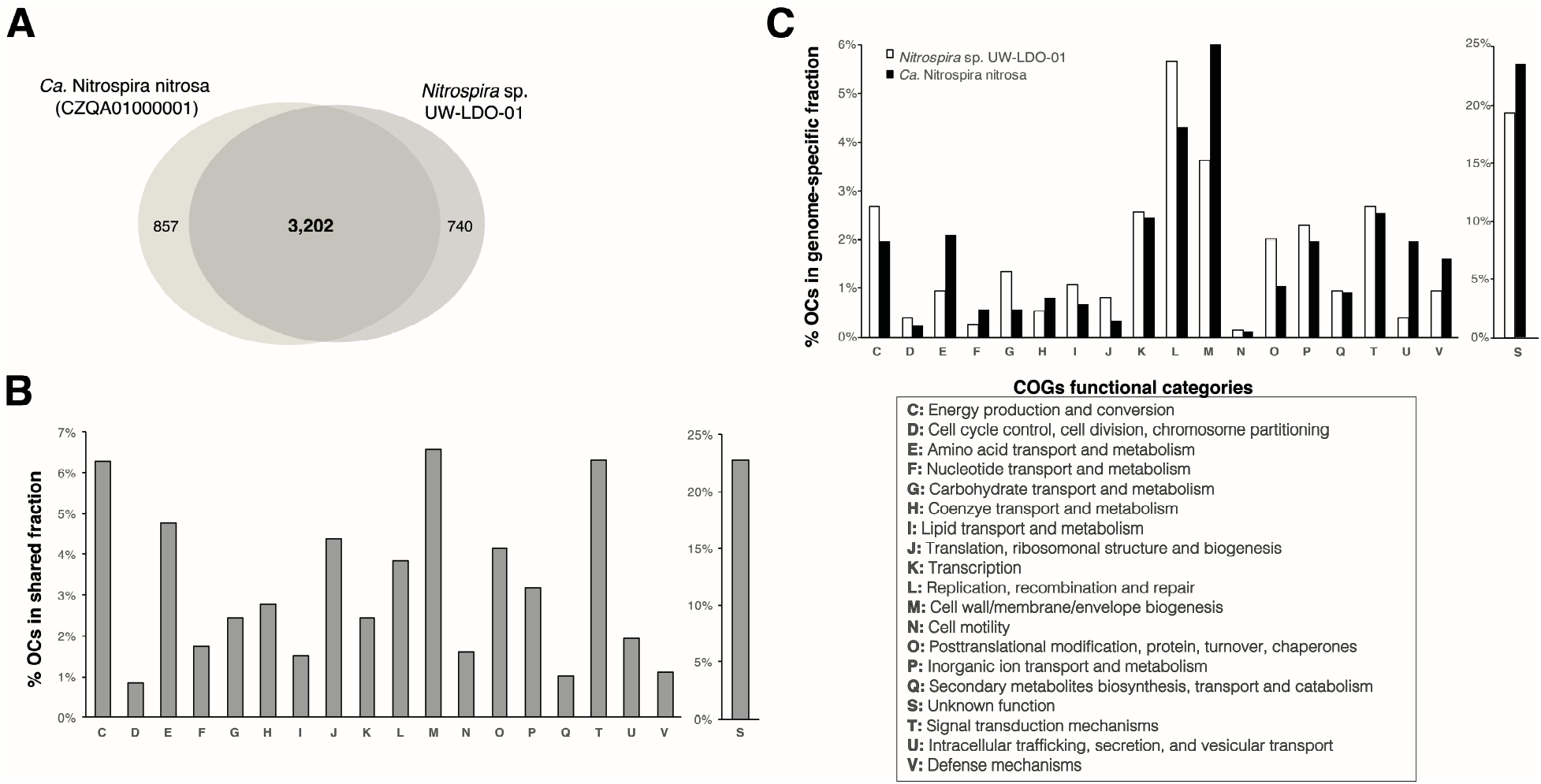
Genomic differences between *Nitrospira sp.* UW-LDO-01 and *Ca.* Nitrospira nitrosa. (A) Venn diagram of ortholog clusters shared between the two draft genomes; (B) distribution of COGs functional classes in the fraction of orthologs shared by the two genomes; (C) distribution of COGs functional classes in ortholog clusters found in only one of the genomes (genome-specific fraction).

OCs belonging to the shared and genome-specific fractions of the two genomes were classified according to their predicted functional role (Fig. 5B-C). 2,906 and 3,009 OCs, in the UW-LDO-01 and *Ca.* Nitrospira nitrosa genomes, respectively, were categorized into functional Clusters of Orthologous Groups (COG) categories. The majority of OCs belonged to the group of “Unknown Function (S)” (23% of shared OCs, and 19% and 23% of the genome-specific fraction in UW-LDO-01 and *Ca.* Nitrospira nitrosa, respectively), indicating a large set of metabolic features not yet elucidated (Fig. 5B-C).

Besides the genes of unknown function, OCs within this shared fraction were mostly represented by the “Cell wall/membrane/envelope biogenesis (M)” (7%), “Signal transduction mechanisms (T)” (6%) and “Energy production and conversion (C)” (6%) families, indicating general conservation of energy metabolism and regulatory mechanisms.

In both genomes, the functional group “Replication, recombination and repair (L)” was overrepresented within the genome-specific fraction. This functional category includes transposases, integrases and other mobile genetic elements, and their extensive representation in the genome-specific fraction indicates that horizontal gene transfer has likely played a significant role in the diversification of *N. nitrosa* strains. The greatest difference among the genome-specific fractions of UW-LDO-01 and *Ca.* Nitrospira nitrosa was the proportion of OCs represented by the “Cell wall/membrane/envelope biogenesis (M)” and “Intracellular trafficking, secretion, and vesicular transport (U)” functional families (Fig. 5C). Glycosyltransferases^35-38^, Type IV^39-41^ secretion system proteins and other enzymes involved in polysaccharide formation (main component of the biofilm matrix) were enriched within these COG categories in the genome-specific fraction of *Ca.* Nitrospira nitrosa. Differences in biofilm formation capabilities between these strains may relate to specific niche adaptation: *Ca.* Nitrospira nitrosa was enriched in a biofilm, whereas UW-LDO-01 was found in a planktonic habitat in wastewater. Analogous findings have been observed in other genera, where differences among biofilm formation capabilities within the same genus were linked to the genome content of different strains.^42-44^ Similar to the results presented here, these genetic differences included the presence of type IV secretion systems and enzymes involved in protein glycosylation.

The comparative genomic analysis also indicated a higher proportion of gene clusters associated with “Lipid transport and metabolism (I)” in *Nitrospira* sp. UW-LDO-01 (Fig. 5C). Genes related to ß-oxidation of long-chain fatty acids to acetyl-CoA were present in the genome of UW-LDO-01, but absent in *Ca.* Nitrospira nitrosa. These genes include a long-chain fatty acid-CoA ligase, acyl-CoA dehydrogenase, enoyl-CoA hydratase, 3-hydroxyacyl-CoA dehydrogenase and acetyl-CoA acetyltransferase. Presence of these lipid-related metabolic genes in other *Nitrospira* strains was confirmed, although the complete pathway is lacking in *Nitrospira defluvii*, *Ca.* Nitrospira nitrificans and *Nitrospira* sp. Ga0074138. This feature may represent a competitive advantage of some *Nitrospira* strains in habitats rich in long-chain fatty acids, such as WWTPs.^45^

### Metabolic Features in *Nitrospira* Genomes

To explore the diverse metabolic capabilities and provide insights into the common and unique metabolic features encoded in the genome of NOB- and comammox-like strains, we compared the gene inventory of 9 complete and draft genomes classified as *Nitrospira*. The analysis was focused on traits associated to energy production, which are summarized in Table S3.

In agreement with previous analyses, only comammox-like genomes encoded ammonia monooxygenase (*amoCAB*) and hydroxylamine dehydrogenase (*haoAB-cycAB*) gene clusters, responsible for ammonia oxidation to nitrite (Table S3), reflecting the capability of this novel *Nitrospira* sub-lineage to perform full-nitrification from ammonia to nitrate.

Analysis of nitrite reducing genes revealed that all *Nitrospira* strains encoded a copper-containing dissimilatory nitrite reductases (*nirK*), which catalyzes the nitrite reduction to nitric oxide, a key step in the denitrification process. Despite the widespread presence of this enzyme across the *Nitrospira* genus, former studies have documented no activity of this protein in NOB-like^46^ or comammox-like strains^1^, where N loss caused by formation of gaseous compounds was not observed. Since it has been predicted that the NXR complex of *Nitrospira* can reduce nitrate to nitrite,^46^ these microorganisms appear genetically capable of converting nitrate (product of nitrification) to nitric oxide. Additional experiments are still needed to obtain more insights into this *Nitrospira* trait. Other denitrification genes, such as nitrate reductase *(nar),* nitric oxide reductase (*nor*) or nitrous oxide reductase (*nos*), were not found in the *Nitrospira* strains analyzed here.

All comammox-like genomes, including UW-LDO-01, encoded the machinery to hydrolyze urea: the *ureABCDFG* urease operon and the *urtABCDE* urea transport system, suggesting that this *Nitrospira* sub-division possesses a high-affinity uptake system for urea and, thus, is adapted to habitats where urea is present at low levels. A gene cluster involved in urea metabolism was also found in *N. moscoviensis* (Table S3), although the urea-binding proteins *urtBCDE* were lacking in the genome. Ureolytic activity of both *N. moscoviensis, Ca.* Nitrospira nitrosa and *Ca.* Nitrospira nitrificans was formerly tested by incubation of these strains with urea-containing media, where urea hydrolysis to ammonium was observed in both cases.^2, 46^ Former studies have also shown the presence of genes for urea utilization in *Nitrospira lenta*,^47^ a novel *Nitrospira* species enriched under low temperatures, suggesting that the ureolytic activity might be associated with this lineage. Since several ammonia oxidizers also possess the capability of hydrolyzing urea,^48-50^ the presence of this trait in *Nitrospira* might evidence horizontal gene transfer among these functional groups.

A contrasting difference among NOB-like and comammox-like genomes was the capability to convert cyanate into ammonia. Only NOB-like genomes encoded a cyanase hydratase enzyme; former studies have experimentally confirmed cyanate degradation in *N. moscoviensis*.^51^ Cyanate is produced intracellularly from urea and carbamoyl phosphate decomposition,^52, 53^ and in the environment from the chemical/physicochemical decomposition of urea or cyanide.^54, 55^ The presence of a cyanase enzyme benefits nitrite oxidizers because it allows them to detoxify cyanate, and the formed ammonium is then available for assimilation and might also serve as a source of energy for ammonia oxidizers in a process described as “reciprocal feeding”.^46, 51^ Further experiments analyzing the effect of cyanate in the growth on comammox-like bacteria are needed to understand how cyanate degradation would confer them a biological advantage, besides generation of ammonia.

The analysis also revealed the presence of the gene inventory for the uptake and oxidation of formate, an exclusive feature of NOB. Growth on formate as electron donor has been confirmed in *N. moscoviensis* (under both micro-oxic and anoxic incubations),^46^ *Nitrospira japonica*^56^ and uncultured *Nitrospira* in activated sludge.^57^ Although formate oxidation should be advantageous for organisms thriving in hypoxic or anoxic habitats, which also includes comammox-like bacteria, this feature is absent in the genome of these microorganisms.

The genome of *N. moscoviensis* encodes a group 2a [Ni-Fe] hydrogenase *(hupS* and *hupL)* and accessory proteins involved in the maturation and transcriptional regulation of hydrogenases *(hypFCDEAB* and *hoxA).* Furthermore, experiments showed that *N. moscoviensis* was capable of growing by aerobic respiration of H_2_.^58^ Although the comammox-like genomes lack the subunits of the [Ni-Fe] hydrogenase (Hup), the five genomes analyzed here encoded a group 3 [Ni-Fe] sulfur-reducing hydrogenase gene set *(hydBGDA and hybD*) positioned at the same locus where Hup is located in *N. moscoviensis*. This hydrogenase complex is an heterotetramer with both hydrogenase activity and sulfur reductase activity, which might play a role in hydrogen cycling during fermentative growth.^59^ Its beta and gamma subunits, that form the sulfur reducing component, catalyze the cytoplasmic production of hydrogen sulfide in the presence of elemental sulfur. The presence of this complex in the genomes indicates the potential of these microorganisms for oxidizing H2 using sulfur as electron acceptor, a trait that has not been analyzed in comammox before, but that could confer this sub-group an advantage when growing under anaerobic conditions. !

Furthermore, the presence of a hyf-like operon (*hyfBCEFGI*), which encodes a putative group 4 hydrogenase complex, was detected in every NOB-like genome, as well as *Ca.* Nitrospira nitrosa, *Nitrospira* sp. Ga0074138 and UW-LDO-01. In *Escherichia coli,* this hydrogenase complex forms part of a second formate hydrogenlyase pathway (oxidation of formate to CO_2_ and reduction of 2H^+^ to H_2_ under fermentative conditions).^60^ This is likely the case for the hydrogenase-4 present in the genome of NOB-like strains, which co-occur with genes encoding formate dehydrogenase. In comammox, however, the role of this distinct hydrogenase is not as clear. In *Ca.* Nitrospira nitrosa, this complex is found immediately adjacent to a carbon monoxide dehydrogenase (CODH), an enzyme that catalyzes the interconversion of CO and CO_2_^61^, a genomic feature that would allow this strain to obtain energy from carbon monoxide.^62^ Conversely, the genomes of *Nitrospira* sp. Ga0074138 and UW-LDO-01 lack the CODH at this position, which in the case of UW-LDO-01, was confirmed by alignment of the metagenomic reads to this gene. No other neighboring gene of the hydrogenase-4 complex could be associated with this enzyme in these two strains, therefore, the biological role of these genes is still unclear.

Altogether, these results reveal specific traits characterizing the NOB and comammox functional groups: while comammox-like *Nitrospira* has the genomic potential of ammonia and nitrite oxidation and potentially, sulfur reduction, NOB-like strains are distinguished by their cyanate degradation and formate oxidation capabilities; and both urea hydrolysis and H_2_ respiration are common traits shared by multiple *Nitrospira* strains.

### The Role of Transcriptional Regulation in *Nitrospira*

Transcriptional regulation of gene expression is the most commonly used strategy to control many of the biological processes in an organism, including progression through the cell cycle, metabolic and physiological balance, and responses to environmental stress. This regulation is generally orchestrated by several transcriptional factors (TFs) that directly coordinate the activity of genes by binding to their promoters. Each *Nitrospira-like* genome codes for at least 100 transcriptional regulators, which account for ~3% of the estimated total number of genes, in agreement with TFs in other microorganisms.^63-65^ A comparative genomic analysis of full and draft *Nitrospira* genomes was used to investigate the repertoire of TFs potentially involved in the survival of these microorganisms under diverse environmental conditions (Table 1).

**Table 1.**
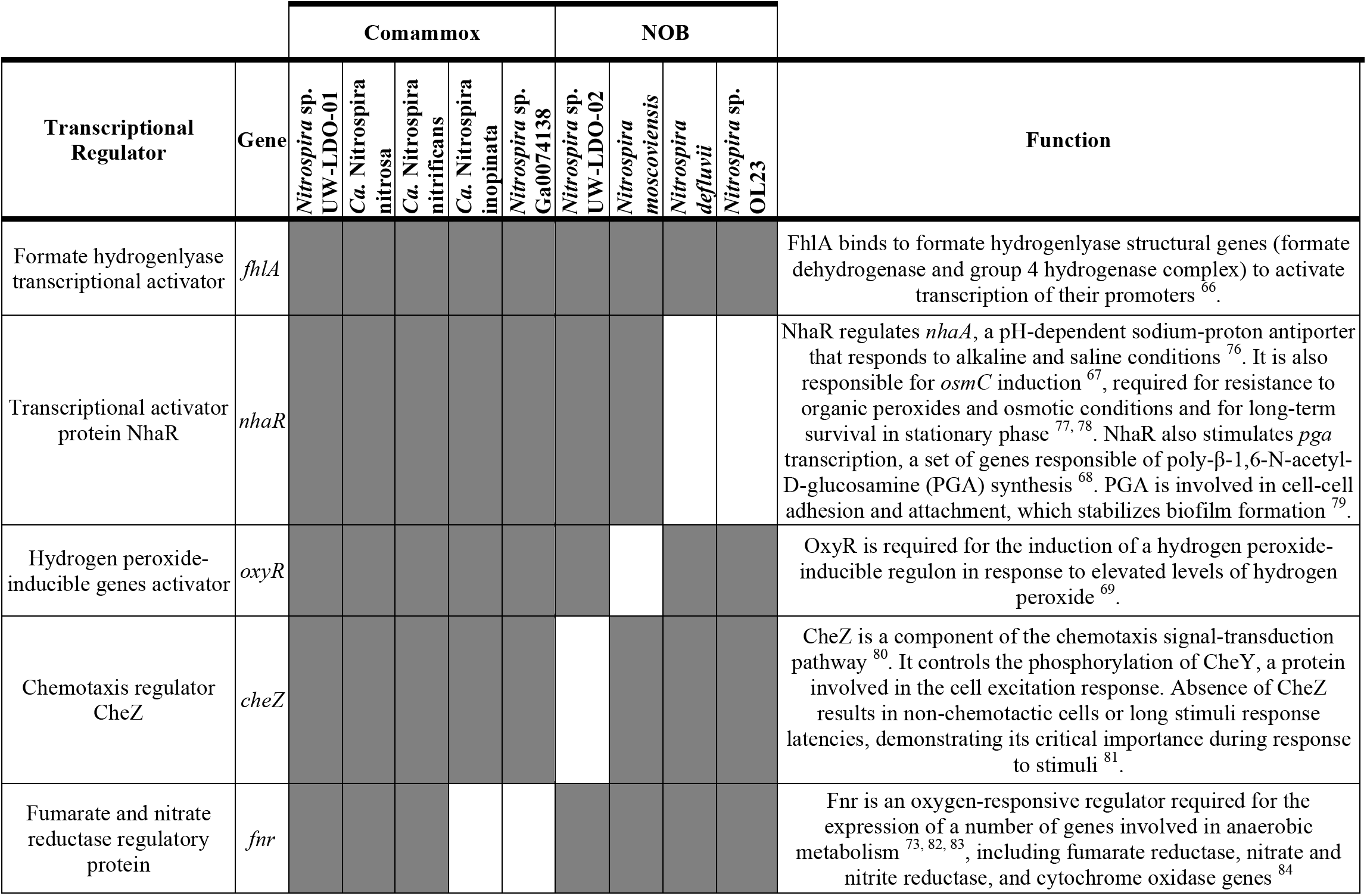
Inventory of transcriptional regulators with implications on adaptive metabolism, from complete and draft genomes of *Nitrospira.* Grey and white rectangles represent presence and absence of each gene, respectively.

Among the TF s analyzed, the formate hydrogenlyase transcriptional activator (fhlA)^60, 66^ was the only one shared across all the *Nitrospira* genomes, although only NOB-like genomes encode genes of its known regulon, the formate hydrogenase complex. The presence of this transcriptional activator in comammox microorganisms, which appear to be genetically incapable of formate oxidation (Table 1), might represent an ancestral trait shared by *Nitrospira* and lost during diversification. This theory would also support the presence of the group 4 hydrogenase (associated with the formate-hydrogenlyase complex in *E. coli)* in both NOB- and comammox-like groups.

A common feature among some NOB and comammox bacteria is the presence of the transcriptional regulators NhaR^67, 68^ and OxyR^69^ (Table 1). The first one is associated with stress response to alkaline, acidic, saline and osmotic conditions. OxyR regulates hydrogen peroxide-inducible genes, such as alkyl hydroperoxide reductase *(ahpCF)* and glutaredoxin *(grxA*), encoded in all the *Nitrospira* genomes. Presence of these genes would confer *Nitrospira* an improved fitness advantage over other nitrifying bacteria. For instance, NhaR is lacking in *Nitrosomonas* and *Nitrobacter* and OxyR is not present in *Nitrosomonas* and *Nitrosospira* (based on genome search). Furthermore, the role of NhaR during regulation of *pga* expression,^68^ allows the biofilm formation process to be considered as a flexible and dynamic developmental process driven by external conditions, representing another means by which NhaR could promote survival of *Nitrospira*. Likewise, the presence of the chemotaxis regulator CheZ in *Nitrospira* suggests chemotaxis as another important mechanism by which these microorganisms efficiently and rapidly respond to changes in the chemical composition of their environment.

To date, the role of the Fnr-type regulatory protein in *Nitrospira* has not been determined. In other microorganisms, Fnr is part of the signaling involved in the adaptation to micro-oxic environments,^70-74^ where it acts as oxygen sensor and regulator of genes involved in anaerobic and micro-aerobic metabolism. In *Nitrospira,* we predict that this TF would regulate similar genes, such as the *frd* operon (fumarate reductase), *sdh* operon (succinate dehydrogenase), *ndh* (NADH dehydrogenase) and *ccb3* complex (cytochrome c oxidase). At least one copy of Fnr in the genomes of UW-LDO-01, *N. moscoviensis, N. defluvii* and N. sp. OLB-3 was located upstream of a copper-containing nitrite reductase gene (nirK), suggesting a possible mechanism that controls expression of this denitrification enzyme. The presence of multiple paralog copies of Fnr in several *Nitrospira* genomes may indicate a rigorous regulation of metabolism when these microorganisms are exposed to low levels of oxygen, an important factor affecting *Nitrospira* community compositions in nitrifying systems.^75^

Overall, this study sheds light about differences in the physiological role of NOB and comammox-like *Nitrospira.* Specifically, the comparative genomic results evidence traits associated with energy metabolism as characteristic to each of these functional groups. Furthermore, the analysis of TFs in *Nitrospira* reveals the alternative use of organic compounds, response to environmental stress, chemotaxis and anaerobic metabolism as some of the key mechanisms for the adaptive metabolism of the genus to multiple and adverse conditions. Further studies in the field should include experiments that combine omics-analysis (transcriptomics, metabolomics, and proteomics) with chemical data to confirm the ecological role and functionality of each of these functional groups and their interactions with other microorganisms.

## ACKNOWLEDGMENTS

This work was partially supported by funding from the National Science Foundation (CBET-1435661 and MCB-1518130) and the Madison Metropolitan Sewerage District. Additional funding from the Chilean National Commission for Scientific and Technological Research (CONICYT) as a fellowship to Pamela Camejo is also acknowledged. The U.S. Environmental Protection Agency, through its Office of Research and Development, partially funded and collaborated in the research described herein. Any opinions expressed in this paper are those of the authors and do not necessarily reflect the views of the agency; therefore, no official endorsement should be inferred. Any mention of trade names or commercial products does not constitute endorsement or recommendation for use.

## ASSOCIATED CONTENT

### Supplementary Material

Supplementary Figures and Tables supporting the information presented in this manuscript.

